# Imprinting and DNA methylation in water lily endosperm: implications for seed evolution

**DOI:** 10.1101/2024.03.15.585284

**Authors:** Rebecca A. Povilus, Caroline A. Martin, Lindsey L. Bechen, Mary Gehring

**Affiliations:** Whitehead Institute for Biomedical Research, 455 Main Street, Cambridge MA 02142; Department of Biology, Massachusetts Institute of Technology, Cambridge MA 02139; Department of Biological Engineering, Massachusetts Institute of Technology, Cambridge MA 02139; Howard Hughes Medical Institute

**Author notes:** Corresponding author: Mary Gehring. **Author Contributions:** R.A.P. and M.G. conceived of the original premise of the study. R.A.P. conducted experiments and wrote the paper with input from M.G. and C.A.M. C.A.M. assisted in processing genomic data. L.L.B performed methylation profiling on leaf tissue. **Competing Interest Statement:** The authors have no competing interests to declare.

**Keywords:** Endosperm, Imprinting, Early Angiosperms, DNA methylation

## Abstract

Endosperm is a key evolutionary innovation associated with the origin of angiosperms (flowering plants). This altruistic seed tissue supports the growth and development of the embryo by mediating the relationship of the mother plant as a nutrient source to the compatriot embryo as a nutrient sink. The endosperm is the primary site of gene imprinting in plants (where expression of an allele in offspring depends on which parent it was inherited from) and of parent-specific epigenetic modifications like DNA methylation, which are differentially patterned during male and female gamete development. Experimental results from a phylogenetically-wide array of monocot and eudicot plants suggest these parent-of-origin effects are a common feature across angiosperms. However, information about genetic imprinting and epigenetic modifications in seeds of angiosperm lineages whose origins predate the monocot-eudicot divergence (such as Nymphaeales, water lilies) is extremely limited. Additionally, Nymphaeales are an intriguing lineage in which to investigate seed genetic and epigenetic phenomena, as they are characterized by diploid endosperm and a maternal storage tissue (perisperm), both of which are unusual across angiosperm diversity. Here, we examined DNA methylation and genetic imprinting using two reproductively compatible water lily sister-species, *Nymphaea thermarum* and *N. dimorpha*. Our results suggest that maternally-expressed imprinted genes and differential DNA methylation of maternally and paternally inherited endosperm genomes are an ancestral condition for endosperm, while other seed characters like seed provisioning strategies, endosperm ploidy, and paternally-expressed imprinted genes might have evolved as coinciding, opposing strategies in the evolutionary dialogue over parental control of offspring development.

## Introduction

The evolutionary origin of endosperm, a second fertilization product in the seeds of flowering plants, fundamentally altered the relationship between an embryo and its mother during seed development. In non-flowering seed plants, the embryo is directly connected to tissue that only contains genome complements from its mother. However, in angiosperm seeds, endosperm largely separates the embryo and its mother and is the product of a fertilization event and thus biparental, with both maternal and paternal genome contributions. Endosperm is widely recognized as a key mediator of developmental and nutritional relationships between an embryo and its mother (Povilus and Gehring 2022). The balance of maternal and paternal genomes is important for endosperm (and seed) viability, as evidenced by the phenomena of parental genome dosage sensitivity ((Haig and Westoby 1991) and references therein). When extra paternal genome complements are added to the endosperm, it over-proliferates – often leading to initially larger but ultimately collapsed, inviable seeds. Conversely, when extra maternal genome complements are added to the endosperm, reduced endosperm proliferation is observed, resulting in smaller seeds with fewer invested maternal resources (Haig and Westoby 1991; Birchler 1993; Scott et al. 1998). The endosperm is subject to other parent-of-origin effects such as imprinted gene expression (where expression of an allele depends on which parent it was inherited from) and parent-of-origin specific epigenetic modifications like DNA and histone methylation, which are differentially patterned during male and female gamete development (Gehring et al. 2006; Park et al. 106; Moreno-Romero et al. 2019; Borg et al. 2020). Our knowledge of endosperm gene imprinting and its underlying mechanisms is built from experiments performed in a phylogenetically wide array of monocot and eudicot plants (Povilus and Gehring 2022; Picard and Gehring 2020). However, information from lineages whose origins predate the monocot-eudicot divergence is extremely limited.

The order Nymphaeales (water lilies) is sister to all other angiosperms except for *Amborella tricopoda*. Endosperm parental genome dosage sensitivity has been documented in a species of water lily (Povilus et al. 2018), but nothing is known about patterns of genetic imprinting or endosperm epigenetic patterning in this or any other ANA-grade lineage (Amborella, Nymphaeales, Austrobaileyales), magnollids, or Chloranthales. In addition to their relationship to other angiosperms, Nymphaeales are a particularly intriguing system in which to investigate genetic imprinting and associated epigenetic patterning given the unique combination of seed characters found in this lineage. First, endosperm of the Nymphaeales is *ab initio*-cellular (the first nuclear division of the endosperm is accompanied by cellular division) and diploid with a 1:1 maternal:paternal genome ratio (Orban and Bouharmont 1998; Williams and Friedman 2002; Friedman 2006; Friedman 2008; Rudall et al. 2008; Povilus et al. 2015), whereas triploid endosperm (2:1 ratio) characterizes the majority of angiosperms and all taxa in which endosperm epigenetic patterning and genetic imprinting have been studied (Haig and Westoby 1991). Diploidy has been suggested to represent the ancestral ploidy of endosperm (Williams and Friedman 2002) and thus Nymphaeales are an opportunity to test how these processes operate in the context of different base maternal-paternal genome/gene dosage ratios. Second, in seeds of Nymphaeales nutrients are primarily stored in a perisperm (which is derived from maternal tissue and contains no paternal genome contribution) instead of in offspring tissues, in contrast to the vast majority of flowering plants (Lersten 2004; Patten et al. 2014). Therefore, Nymphaeales is an excellent clade in which to investigate the suggested connection between genetic imprinting in endosperm and control of nutrient storage (Haig and Westoby 1991; Patten 2014). Nutrient storage in perisperm is only initiated after fertilization (Povilus et al. 2015), suggesting influence of offspring tissues on this process.

Characterizing genetic imprinting and epigenetic modifications in water lilies therefore offers a unique perspective on the evolution of key endosperm traits and processes that are associated with the origin of angiosperms. Here, we sought to determine whether gene imprinting and parent-of-origin effects on DNA methylation, which has been mechanistically linked to gene imprinting, preceded the origin of parental dosage-imbalanced (triploid) endosperm.

## Results

*Nymphaea thermarum* has been developed as an experimental system for the Nymphaeales (Povilus et al. 2015; Povilus et al. 2018; Povilus et al. 2020). Assessing parent-of-origin effects at the molecular level requires sequence polymorphisms, of which there are few within the highly-inbred extant populations of *N. thermarum* in cultivation (Povilus et al. 2015). We therefore assessed parent-of-origin effects in Nymphaeales by examining F1 tissue from crosses between *N. thermarum* and *Nymphaea dimorpha* (which was formerly known as *N. minuta*). These two species are estimated to have diverged roughly 20 million years ago (Borsch et al. 2011). We confirmed the internal structure of young seeds of *N. thermarum*, *N. dimorpha*, and of F1 reciprocal crosses and determined that we could ensure consistency in developmental stage among crosses (Figure 1A). The hybrid F1 seeds are fully viable and germinate to give rise to viable F1 plants (Supplementary Figure 1), suggesting no large-scale divergence in endosperm developmental programs that would lead to failure in seed development. We performed long-read based, *de novo* genome assembly and annotation for *N. dimorpha* (248 contigs with an NG50 of 13,941,033 bp, representing 83% of the estimated genome size, with the set of 40,850 annotated genes having a BUSCO score of 85% for the Embryophyta gene set) and an improved genome assembly and annotation for *N. thermarum* (1,553 contigs with and NG50 of 4,352,861, representing 86% of the estimated genome size, with the set of 42,431 annotated genes having a BUSCO score of 83% for the Embryophyta gene set) (Supplementary Table 1, Supplementary Figure 2). To allow direct comparisons of genomic regions, the *N. thermarum* and *N. dimorpha* genomes were aligned and re-annotated to create “reorganized” genomes for each species (each 358,929,111 bp in length and with a resolved annotation having 39,608 genes and a BUSCO score of 72% for the Embryophyta gene set).

**Figure 1:**
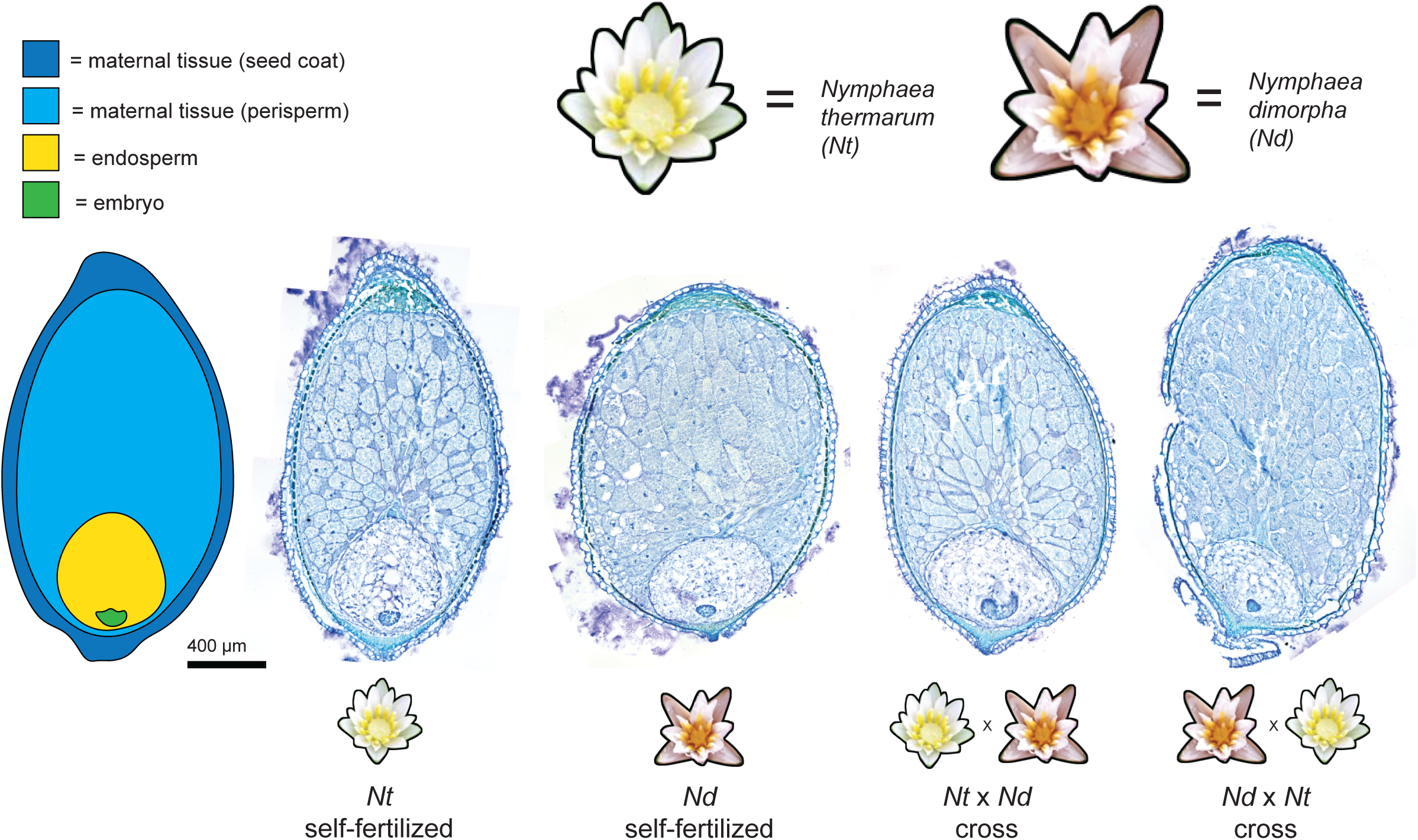
Seed structure in water lilies and F1 hybrids. Young seeds (9-10 days after anthesis) of self-fertilized *N. thermarum* (*Nt*) and *N. dimorpha* (*Nd*), as well as reciprocal crosses between the two species. In all seeds, young embryos are surrounded by cellular, diploid endosperm, which in turn is surrounded by a maternal nutrient storage tissue (perisperm).

To examine parent-of-origin specific gene expression, we made use of reciprocal crosses between *N. thermarum* (*Nt*) and *N. dimorpha* (*Nd*) (2 samples each from *N. thermarum x N. dimorpha* and *N. dimorpha x N. thermarum* crosses) as well as self-fertilized seeds (3 samples each of *N. thermarum* and *N. dimorpha*), and isolated RNA from young endosperm at 9-10 days after pollination/anthesis (Supplementary Table 1). By performing mRNA-seq, we detected expression of a total of 22,984 genes with a TPM >= 1 (expression averaged across all samples). A principal component analysis (PCA) of total gene expression revealed that biological replicates clustered together according to cross type, with hybrid endosperm samples midway along PC1 (54.85%) between endosperm from *N. thermarum* self-fertilized seeds and *N. dimorpha* self-fertilized seeds (Figure 2A). Differential gene expression analysis between sets of hybrid and non-hybrid seeds revealed that 879 genes were consistently significantly differentially expressed in all comparisons; this set of genes was not significantly enriched for any KEGG pathways, but was enriched for the GO biological process term “RNA-dependent DNA biosynthetic processes” (FDR = 1.9e-2, n=8, fold enrichment = 7.1). Importantly, while expression differences between hybrid and non-hybrid endosperm existed, endosperm of the two hybrid cross directions were more similar to each other than they were to endosperm of the parental lines (Figure 2A). We implemented a previously developed pipeline to evaluate imprinted gene expression (Gehring et al, 2011) (see Methods). In each possible comparison of a *Nt* x *Nd* and *Nd* x *Nt* cross, we identified transcripts that showed a significant bias in the number of reads mapping uniquely to either the maternally-or paternally-inherited alleles, in both cross directions (Figure 2B, Supplementary Table 2). For these imprinting tests, 26,465 genes had at least one read that could be assigned to a parent-of-origin and 16,647 genes passed our minimum allele-specific read count cut-off of 50 reads and were assessed for imprinting. Our analysis revealed the presence of imprinted genes in *Nymphaea*. We identified small numbers of paternally-biased genes in individual cross comparisons, but only 1 paternally expressed imprinted gene (PEG) was consistent in at least 75% of comparisons (3 of 4 total possible cross comparisons) (Figure 2C); this PEG is a homolog of *CELLULOSE SYNTHASE LIKE G2* (*ATCLSG2*) in *Arabidopsis thaliana*. A handful of PEGs have previously been identified as conserved between monocots and dicots (Pignatta and Gehring 2012). We examined the expression of homologs of these specific genes in *Nymphaea* endosperm. Although there was some evidence for paternally-biased expression, there were also large *cis* or species effects on transcription, and these genes did not meet all of our criteria for imprinting (Supplementary Table 2). In contrast to PEGs, 157 MEGs were consistently identified in at least 75% of comparisons, with 147 being identified as MEGs in all comparisons (Figure 2C) (Supplementary Table 2). Previous studies have shown that imprinting can be altered or obscured in interspecies hybrids (Josefsson et al. 2006; Burkart-Waco et al. 2015; Florez-Rueda et al. 2016), and we evaluated whether the identified MEGs were differentially expressed in the F1 hybrid endosperm. When endosperm gene expression profiles were compared between each hybrid type and each parental species type (Figure 2D, Supplementary Figure 3, Supplementary Table 3), only 10 MEGs were consistently (in 75% or more of comparisons) significantly differentially expressed, including only 1 MEG that was significantly differentially expressed in all comparisons. This indicates low overlap between identified MEGs and genes that are mis-regulated in hybrid crosses. We conclude that imprinted expression exists in *Nymphaea* endosperm, but is largely restricted to MEGs and mostly does not include genes whose expression is altered in hybrid endosperm.

**Figure 2:**
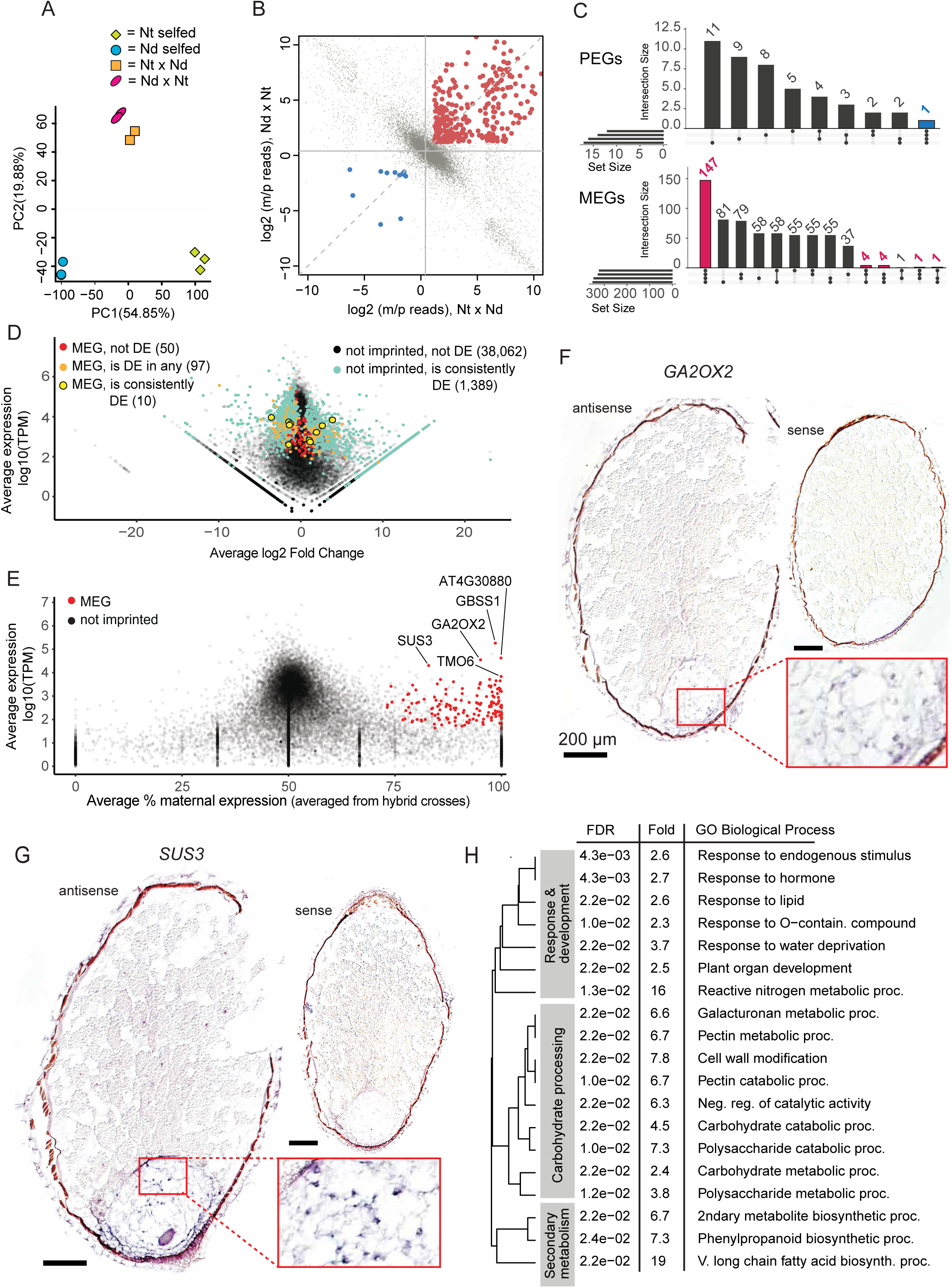
Genetic imprinting in water lily endosperm. A) Principal component analysis of RNA-seq data from young endosperm samples isolated from self-fertilized *N. thermaurm* (Nt) and *N. dimorpha* (Nd) seeds, and reciprocal crosses of *N. thermaurm* and *N. dimorpha*. B) Plot of the ratio of maternal to paternal allele reads in endosperm of one reciprocal cross comparison. Female in cross listed first. Genes that pass the stringency cut-offs for being called as imprinted are highlighted in red (MEGs, maternally expressed genes) or blue (PEGs, paternally expressed genes). Gray dots indicate genes that are not called as imprinted. C) Upset plot showing the number and consistency of genes called as MEGs or PEGs across comparisons of different replicates. MEGs and PEGs called in at least 75% of comparisons are highlighted in blue (PEGs) or red (MEGs). D) Average expression and average log2 fold change of genes expressed > 1 TPM, in the comparison of Nt self-fertilized and Nt x Nd hybrid endosperm. Genes are color-coded according to whether they are significantly differentially expressed in no comparisons or consistently (75% or more of comparisons), and/or were identified as a MEG in no samples, in any samples, or consistently (75% or more of samples). Number of genes in each category is noted. Similar graphs for individual comparison types are shown in Supplementary Figure 3. E) Average expression and average percent maternal expression for all expressed genes (TPM > 1). Genes called as MEGs in at least 75% of replicate comparisons are shown in red, with putative *Arabidopsis thaliana* homology indicated for some MEGs of interest. F) In situ hybridization of putative homolog of *GA2OX2*. Inset shows magnification of endosperm of sample treated with antisense probe. Scalebars = 200 μm. G) In situ hybridization of putative homolog of *SUS3*. Inset shows magnification of endosperm of sample treated with antisense probe. Scalebars = 200 μm. H) GO-enrichment analysis (for biological process terms) of genes consistently called as MEG in the endosperm, reported with FDR-adjusted p-value and fold-enrichment.

The identification of MEGs can be influenced by contamination with maternal tissue or transcripts in endosperm samples and thus we evaluated the extent to which this might be affecting our results. Notably, PCA results indicate that hybrid endosperm from both cross directions was more like each other transcriptionally than endosperm of their mother species, suggesting no significant, wide-spread maternal contamination (Figure 2A). We additionally performed RNA *in situ* hybridizations on seeds of self-fertilized *N. thermarum* for a set of identified MEGs to test whether they showed substantial expression in maternal tissue or lacked expression in the endosperm (both of which would indicate potential for maternal tissue contamination). Target genes were selected for high maternal expression bias and high expression (Figure 2E), including a gibberellic acid oxidase homolog (*GA2OX2*) (Figure 2F) and a sucrose synthase 3 homolog (*SUS3*) (Figure 2G). We furthermore confirmed that the high starch and carbohydrate content of the perisperm was not interfering with the *in situ* hybridization experimental protocol, as we were able to detect expression of a subfamily of terpene synthases in the perisperm, as well as in the endosperm (Supplementary Figure 4). For both *GA2OX2* and *SUS3*, we detected expression in the endosperm and not in the perisperm, indicating that for these and likely other MEGs, the identified maternal expression bias is unlikely to be caused by contamination with maternal tissue. For *SUS3*, we also performed *in situ* hybridizations on seeds from reciprocal crosses and from *N. dimorpha* self-fertilizations, and found similar endosperm expression patterns (Supplementary Figure 4).

Additionally, we took advantage of whole-seed expression datasets from *N. thermarum* (Povilus and Friedman 2022) to explore the potential impact of maternal tissue transcript contamination on our imprinting results. For each transcript expressed in the endosperm, we calculated corrected endosperm maternal read counts based on assuming that 50% or 25% of the isolated endosperm transcript pool was comprised of transcripts from a whole-seed pool (Tonosaki et al. 2024). We found that correcting for an assumed 50% sample contamination resulted in identification of 112 MEGs and correcting for an assumed 25% contamination resulted in identification of 134 MEGs, compared to the originally identified 157 MEGs (Supplementary Table 4). Both sets of 112 MEGs and 134 MEGs were subsets of the originally identified 157 MEGs. The “corrected” MEGs were enriched for KEGG or GO biological process terms related to response and development, carbohydrate processing, and secondary metabolism (Supplementary Table 4), similar to the enrichments for the original, uncorrected set of MEGs (Figure 2G). Furthermore, in the corrected datasets *SUS3* was not identified as a MEG, although our *in situ* hybridizations demonstrated that *SUS3* was not highly expressed in maternal seed tissues but was expressed in endosperm (Figure 2F, Supplementary Figure 4). We concluded that the correction for extensive, assumed whole-seed transcript contamination was likely inappropriately removing true MEGs, and therefore proceeded to use the uncorrected set of MEGs for further analysis.

Overall, *Nymphaea* MEGs were significantly enriched for GO annotations associated with response and development, carbohydrate processing, and secondary metabolism (Figure 2H, Supplementary Table 4). The enrichment for processes integral to development and nutrient dynamics in seeds is similar to what has been found in other species (Xin et a. 2013; Picard et al. 2021). Notably, *SUS3* is a maternally expressed imprinted gene in *Nymphaea* and is a key part of nutrient processing in the endosperm in other species (Angeles-Núñez and Tiessen 2010). These results are congruent with genetic imprinting being associated with nutrient dynamics during seed development.

Having found evidence for parent-of-origin effects on gene expression, we next investigated parent-of-origin effects on DNA methylation by performing endosperm whole-genome enzymatic methyl-sequencing (EM-seq). We again made use of reciprocal crosses between *N. thermarum* and *N. dimorpha* to permit allele-specific characterization of DNA methylation patterning in young F1 hybrid endosperm; biological replicates from two *N. thermarum x N. dimorpha* crosses and from three *N. dimorpha x N. thermarum* crosses were analyzed (Supplementary Table 1). We also obtained methylation profiles from single samples of *N. thermarum* and *N. dimorpha* leaves and the leaves of an F1 *N. dimorpha x N. thermarum* hybrid. In both endosperm and leaves, the average methylation profiles of *N. thermarum* and *N. dimorpha* alleles of genes and repeats (transposable elements) (Figure 3A) was similar to other angiosperms (Niederhuth et al. 2016), with CG methylation occurring in gene bodies and CG and non-CG methylation in repeats. We identified differentially methylated regions (DMRs) between *N. thermarum* and *N. dimporpha* genomes in F1 hybrid leaves and between *N. thermarum* and *N. dimorpha* leaves. In hybrid leaves, the majority of DMRs occurred in the CG context, and similar numbers of regions were more methylated in one species or one genome versus the other (Figure 3B, Supplementary Figure 5). We then identified DMRs between maternal and paternal alleles in F1 endosperm. The majority of CG and CHH DMRs were less methylated on maternal alleles than on paternal alleles, regardless of which species was the maternal parent in the F1 hybrid endosperm (Figure 3B). Additionally, maternal alleles of both species were consistently hypermethylated in the CHG context and were generally more methylated in endosperm than in hybrid or parental species’ leaf tissue, for both genic flanking regions and within repeats (Figure 3A). The higher methylation of maternal alleles in the CHG context was also borne out by analyzing the number of DMRs between maternal and paternal alleles (Figure 3B, Supplementary Table 5): the CHG context had the highest difference between the portion of maternally-or paternally-hypermethylated DMR windows, with both species having more maternally hypermethylated windows. These methylation patterns were not found in leaf tissue (Figure 3B; Supplementary Figure 5) and may therefore be unique to endosperm. Together, these findings parallel observations of methylation patterning in endosperm of monocots and dicots, in which endosperm is CG hypomethylated on maternal alleles due to active DNA demethylation that occurs in the central cell (the female gamete that is the progenitor of the endosperm) before fertilization. Maternal allele CG hypomethylation has been noted in rice, Arabidopsis, and other species (Gehring et al. 2009; Hsieh et al. 2009; Waters et al. 2011; Rodrigues et al. 2013; Park et al. 2016; Zhang et al. 2016; Zhang et al. 2021). Endosperm maternal allele CHG hypermethylation has been observed in *Arabidopsis lyrata* (Klosinska et al. 2016) and to a lesser extent in *A. thaliana* (Moreno-Romero et al. 2019), although this occurs primarily in gene bodies, unlike the observations here where it occurs in gene flanking regions and in repeats.

**Figure 3:**
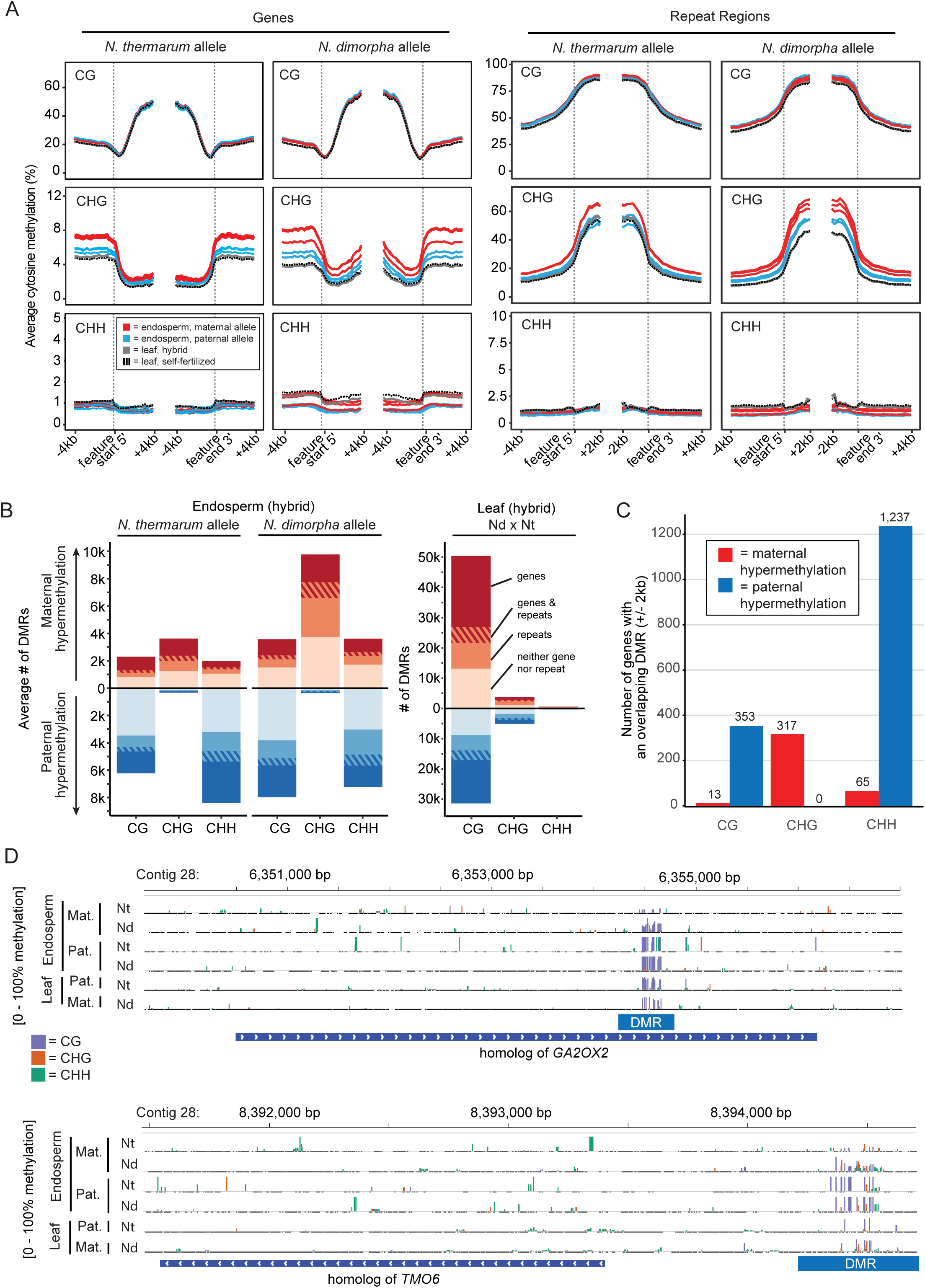
DNA methylation in water lily endosperm. A) Average parental allele cytosine methylation across genes or repeats in young endosperm of reciprocal crosses between *N. thermarum* and *N. dimorpha*, aligned at either the 5’ transcription start site or 3’ transcription end site. Each track represents the maternal or paternal alleles of an individual sampled cross (N=3 for *N. thermarum* x *N. dimorpha* crosses, N = 2 for *N. dimorpha* x *N. thermarum* crosses). Tracks are grouped by species and color-coded to indicate whether the track represents the maternally-or paternally-inherited alleles. B) Average number of maternally- or paternally-hypermethylated DMRs that overlap a gene and/or a repeat region, for each species. DMRs were called by comparing the genomes of each species when they were maternally- or paternally-inherited , for hybrid endosperm and leaf tissue (only one cross direction performed for leaf tissue). Red bars (above 0) indicate maternal hypermethylation, blue bars (below 0) indicate paternal hypermethylation. Bars are further color-coded to indicate the proportion of DMRs that overlap a gene, a gene and repeat, a repeat, or neither gene nor repeat. C) Number of genes (+/-2kb) that consistently overlap maternally- or paternally-hypermethylated DMRs (gene had to have at least one DMR overlap of the indicated type in at least 75% of comparisons, as well as have an overlapping DMR of the opposite type in no more than 25% of comparisons). D) Genome browser snapshots of DNA methylation for homologs of *GA2OX2* and *TMO6*, showing examples of methylation patterning (blue = CG, orange = CHG, and green = CHH) on maternal or paternal genomes, for each species as the maternal and paternal genome, in endosperm and leaf tissue.

We further examined the set of genes that overlapped a maternal-or-paternal hypermethylated DMR consistently across both species (had a DMR that overlapped the gene region +/-2kb in at least 75% of pairwise comparisons, while not overlapping a DMR of the opposite type in more than 25% of comparisons) (Figure 3C, Supplementary Table 5). Similar to looking at numbers of DMRs associated with genes separately in each species, more genes were associated with paternally hypermethylated DMRs in the CG and CHH contexts, while in the CHG context more genes overlapped with maternally hypermethylated DMRs. Few imprinted loci were consistently associated with allele-specific DMRs (Supplementary Table 5). Four MEGs were consistently associated with maternal allele hypomethylated DMRs in the CG context, including homologs of *TARGET OF MONOPTEROS 6* (*TMO6*) and *GA2OX2* (Figure 3D), and 2 MEGs were associated with maternally hypomethylated DMRs in the CHH context. These regions were not differentially methylated in F1 hybrid leaves, indicating a parent-of-origin effect on methylation that is specific to the endosperm (Figure 3D). The one identified PEG was not associated with any significant methylation differences between parental alleles. Thus, we conclude that there are parent-of-origin specific differences in DNA methylation in *Nymphaea* endosperm, of a similar nature (maternal CG hypomethylation) to those observed in monocots and eudicots. The majority of *Nymphaea* MEGs are not associated with differential DNA methylation. For comparison, in maize and Arabidopsis approximately 50% and 40% of MEGs, respectively, are associated with differential DNA methylation (Pignatta et al. 2014; Gent et al. 2022).

## Discussion

Our results illuminate the evolution of imprinting and potential mechanisms facilitating the emergence of gene imprinting. In summary, we found that genetic imprinting and parent-of-origin effects on DNA methylation patterning occur in endosperm of Nymphaea seeds. Both DNA methylation and genetic imprinting have been suggested to be strategies that can alter the effective maternal or paternal genome/gene dosage in endosperm. Changes to absolute parental genome dosage (and ploidy) of endosperm have also occurred repeatedly during angiosperm evolution. Our discovery of endosperm genetic imprinting and parent-of-origin effects on DNA methylation in *Nymphaea* suggests that these characters/processes predate the evolution of triploid endosperm and are likely to be have been either co-opted from preexisting ancestral molecular programs or are novelties associated with the origin of endosperm itself (Figure 4). In either case, these findings demonstrate that a 2:1 maternal:paternal genome dosage ratio is not a requirement for either endosperm maternal allele CG hypomethylation and CHG hypermethylation, or for genetic imprinting. Furthermore, our results suggest that parent-of-origin effects on endosperm development in *Nymphaea* (Povilus et al. 2018) could be linked to parent-of-origin-specific DNA methylation patterning or maternally-expressed imprinted genes, but not extensively to paternally-expressed imprinted genes. This is perhaps surprising given that Povilus et al. (2018) observed paternal effects when diploids were pollinated by tetraploids: in mature seeds, endosperm of both 4n x 2n (maternal excess) and 2n x 4n (paternal excess) is larger than endosperm of 2n x 2n crosses. However, the developmental timing by which larger endosperm is achieved differs between maternal and paternal excess crosses. During later development (7-32 DAA), the endosperm of 2n x 4n crosses grows significantly faster than that of 2n x 2n crosses. By contrast, in 4n x 2n crosses, the endosperm grows faster earlier (1-7 DAA), and then decelerates at later stages. The observed maternal and paternal effects in *N. thermarum* endosperm are therefore distinct from those typically observed in maize or Arabidopsis, where maternal excess seeds undergo early endosperm cellularization and are typically smaller than 2n x 2n seed at maturity and paternal excess seeds undergo extended endosperm proliferation and are larger and dead (Arabidopsis) or smaller and dead (maize) at maturity (Scott et al. 1998; Pennington et al. 2008). Although it has been proposed that an increased dosage of PEGs is the cause of interploidy paternal excess phenotypes, direct evidence is limited. Indeed, in Arabidopsis it has been shown that PEG expression is increased in both viable and non-viable seeds from Arabidopsis paternal excess crosses (Satyaki and Gehring 2019), suggesting that PEG expression is not the determining factor, or sole determining factor, of paternal excess interploidy phenotypes. Finally, a single asymmetry between parental genomes, such as a MEG, has the potential to cause both maternal and paternal parental effects as the endosperm dosage of the gene would differ between interploidy crosses.

**Figure 4:**
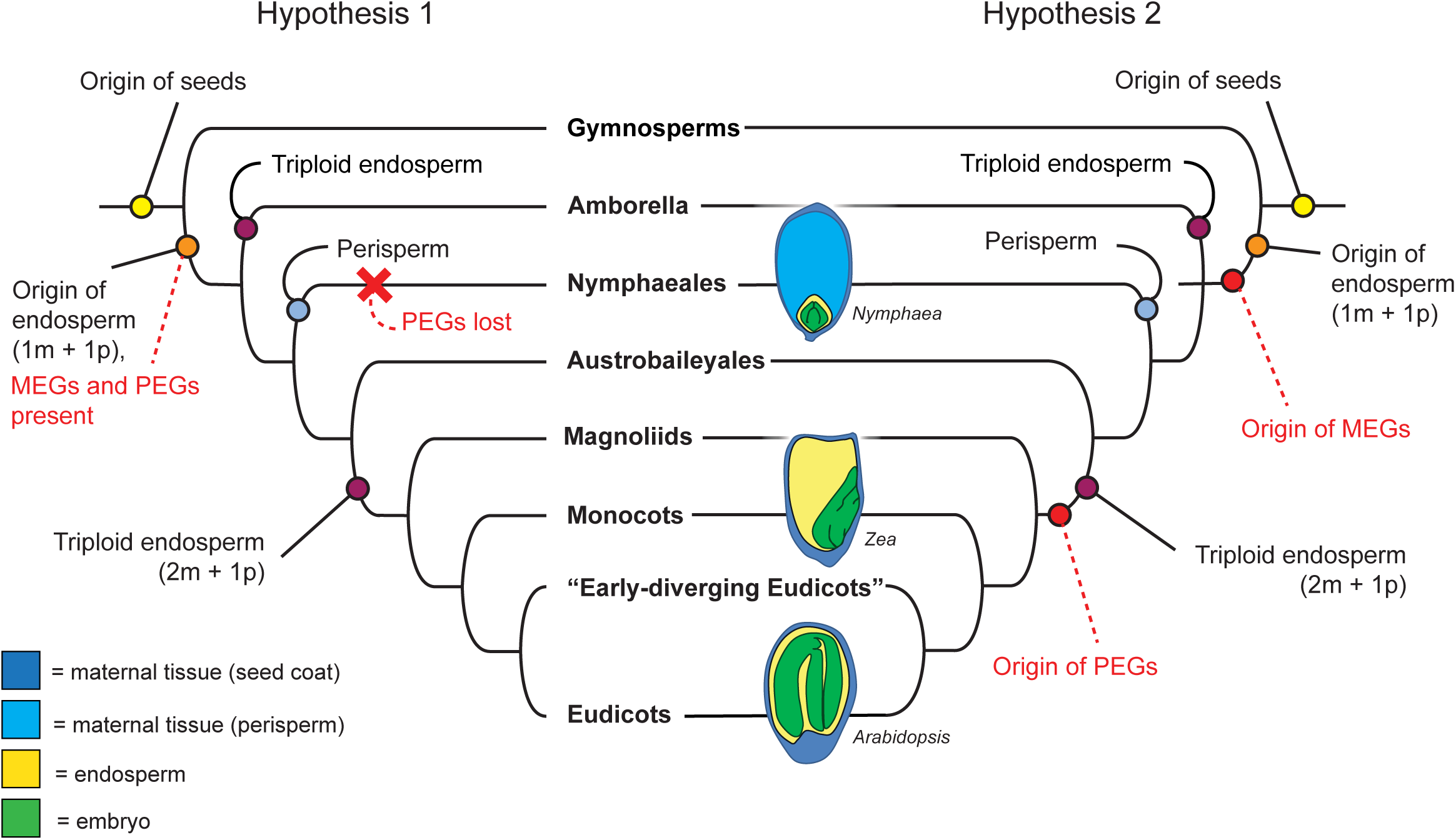
Two hypotheses regarding the evolution of genetic imprinting in endosperm, endosperm ploidy, and nutrient storage strategies in angiosperm seeds. Hypothesis 1 posits that MEGs and PEGs were originally present in endosperm of the last common ancestor of angiosperms, and that PEGs were lost in the water lily lineage in association with the origin of perisperm. Hypothesis 2 posits that MEGs evolved as a maternal response to the addition of a paternal genome complement in a nutrient-mediating tissue, while PEGs evolved as a response to the origin of triploid endosperm, which features the addition of a second maternal genome complement. Dashed lines indicate events hypothesized as a result of this study. Seed diagrams show diversity of mature seed structures, highlighting diversity in developmental origin of the primary site of nutrient storage (*Nymphaea* = maternal perisperm; *Zea* = endosperm; *Arabidopsis* = embryo).

It remains possible that we failed to detect PEGs because of our use of inter-species comparisons or because of the developmental stage at which we analyzed imprinting. Inter-specific and inter-ploidy crosses can result in misregulation of imprinted gene expression. In hybrid crosses between *A. thaliana* and *A. arenosa*, several PEGs gain expression from the maternally-inherited allele in whole seed data, rendering them biallelic or even maternally biased (Josefsson et al. 2006; Burkart-Waco et al. 2015). Other genes become newly paternally biased. Similarly, endosperm of wild tomato hybrids exhibit a genome-wide increase in maternal transcriptome contribution and reduced paternal bias of PEGs (Florez-Rueda et al. 2016). However, in both of these examples, endosperm development is defective and the hybrid seeds are inviable. This is not the case in crosses between *N. thermarum* and *N. dimorpha* – the seeds and F1 plants are fully viable (Figure 1, Supplementary Figure 1), suggesting no defects in endosperm development of the kind that is often correlated with misregulated genomic dosage or misregulated imprinting. Thus, although we cannot exclude it, we think it unlikely that PEGs were not detected in *Nymphaea* endosperm because of the genetic material used in our crosses. We also cannot exclude the possibility that PEGs might be active at earlier stages of Nymphaea seed development than we sampled, before significant development/functioning of the perisperm, as has been suggested by a study on endosperm of a different Nymphaea species (Florez-Rueda et al, 2024). Yet, in other species PEGs have been detected across all assayed stages of endosperm development, including in seeds with similar stages of embryo development as sampled in our study.

The observed differences in CG DNA methylation between endosperm maternal and paternal alleles (Figure 3B,D) are consistent with the activity of a 5-methylcytosine DNA glycosylase in the central cell before fertilization, which would be predicted to cause maternal allele CG hypomethylation in endosperm after fertilization. Homologues of DME are present in all angiosperms (Pei et al. 2019), including *Nymphaea* (Povilus and Friedman 2022). The lack of high congruence between CG maternal allele hypomethylated DMRs and *Nymphaea* MEGs is not inconsistent with data from other species. In Arabidopsis and maize, there are many more endosperm CG hypomethylated regions than there are imprinted genes (Gehring et al. 2009; Pignatta et al. 2014; Gent et al. 2022). Although the imprinting of individual Arabidopsis MEGs like *FWA* and *SDC* is correlated with differential methylation between maternal and paternal alleles (Kinoshita et al. 2004), as a group MEGs are not enriched for differential methylation compared to non-imprinted genes (Pignatta et al. 2014). This is in contrast to PEGs, where differential CG methylation of upstream or downstream regions is enriched compared to non-imprinted genes (Pignatta et al. 2014; Moreno-Romero et al. 2019). We also observed endosperm maternal allele CHG hypermethylation in *Nymphaea* endosperm in repeats and gene-flanking regions (Figure 3A-C). In other angiosperm species, CHG methylation is frequently associated with H3K9me2 and transcriptional silencing. The maternal alleles of PEGs in Arabidopsis species are significantly associated with CG hypomethylation and CHG hypermethylation in endosperm (Klosinska et al. 2016; Moreno-Romero et al. 2019). Although we observed maternal allele CG hypomethylation and CHG hypermethylation in *Nymphaea* endosperm, they were not associated with PEGs, suggesting that paternal expression bias is not an inevitable outcome of these types of epigenetic patterns. Other epigenetic mechanisms could regulate gene imprinting in *Nymphaea*. In mammals, imprinting of a small number of genes is regulated by H3K27me3, without an apparent role for DNA methylation (Lewis et al. 2004; Inoue et al. 2017; Santini et al. 2021; Raas et al. 2022), and has been proposed as an ancestral mechanism of imprinting in the placenta. H3K27me3 also plays an important role in plant gene imprinting and is often coincident with differential DNA methylation (Gehring et al. 2006; Zhang et al. 2014; Moreno-Romero et al. 2019). Histone modification patterns in *Nymphaea* endosperm is a potential area of future investigation.

In the context of parental conflict over the investment of maternal resources in offspring and maternal-offspring coadaptation, the evolution of maternal storage tissues and the notable paucity of PEGs in Nymphaea endosperm give rise to two distinct hypotheses about the early evolution of angiosperm seed development and endosperm molecular/genetic processes (Figure 4). For the first hypothesis, if MEGs and PEGs were both present in the ancestrally diploid endosperm of the earliest flowering plants, then PEGs were largely lost in association with the evolution of perisperm in the Nymphaeales (Figure 4, left). If genetic imprinting is a tool to establish maternal or paternal control over resource investment in offspring (Haig and Westoby 1991), perhaps PEGs are no longer an effective paternal strategy when the mother establishes primary control by storing invested resources in a maternally-derived tissue that does not have a paternal genome contribution. A second hypothesis is that genetic imprinting has evolved in stepwise fashion along with endosperm ploidy changes (Figure 4, right). In this case, MEGs may have evolved as a maternal strategy to balance the addition of a paternal genome – and potential for paternal influence on seed development – that resulted in the origin of endosperm. Subsequent addition of a second maternal genome complement with the evolution of triploid endosperm may then be similarly associated with the evolution of PEGs. Studying other members of ANA-grade lineages with diploid endosperm that lack perisperm (such as in Austrobaileyales (Losada et al. 2017)) or taxa with triploid endosperm and perisperm (as can be found in Amaranthaceae (Coimbra and Salema 1999; López-Fernández and Maldonado 2013), within the eudicots) would help distinguish between these hypotheses by specifically testing the relationship between maternally-derived storage tissues and the absence of PEGs. Thus, while characterizing parent-of-origin effects on gene expression and epigenetic modifications in Nymphaea endosperm is an important step in understanding the evolution of molecular processes in endosperm, the genetic and developmental diversity across angiosperm seeds deserves further attention.

## Materials and Methods

### Data availability

Data generated as part of this study are available as part of NCBI BioProjects PRJNA1085993, PRJNA1086866, PRJNA1086863, PRJNA1085992, and PRJNA1087317, including raw sequence data deposited in SRA, and genome assemblies deposited in NCBI WGS.

### Plant growth and sample collection

Seeds of *N. thermarum* and *N. dimorpha* were sourced from the Arnold Arboretum of Harvard University (Boston, MA, USA) and grown at the Whitehead Institute for Biomedical Research (Cambridge, MA, USA) (Supplementary Materials and Methods).

Controlled pollinations and self-fertilizations were performed as previously described (Povilus et al. 2015; Povilus et al. 2018). For collection of seeds from crosses and self-fertilizations, first day of anthesis (time of female receptivity and fertilization (Povilus et al. 2015)) was defined by presence of stigmatic fluid. Fruits were collected at 10 days after anthesis (DAA) and seeds were immediately removed and dissected with fine forceps in dissection buffer. Endosperm tissue was washed with dissection buffer multiple times and frozen in liquid nitrogen (Supplementary Materials and Methods).

### Whole genome sequencing, assembly, and annotation

For long-read and short-read DNA sequencing, high molecular weight DNA was extracted from > 1 g young leaf samples from a single individual plant using a modified CTAB-based protocol (Supplementary Materials and Methods). Samples were prepared for PacBio sequencing (PacBio Sequel SMRTcell, 20h, v3 chemistry) and were sequenced at the MIT BioMicroCenter. *Nymphaea thermarum* was sequenced in both LR (long-read) and HiFi (high-fidelity) modes; *N. dimorpha* was sequenced in LR mode. *N. dimorpha* short-read data was obtained from the same sample, using 1 lane of an Illumina HiSeq2000 flow-cell (40 bp, paired-end reads) at the MIT BioMicroCenter. Short-read genomic data for *N. thermarum* was downloaded from BioProject PRJNA508901.

Genome assembly for *N. thermarum* and *N. dimorpha* was performed separately using long-reads as input for Canu (version 2.1.1) (Koren et al. 2017); short-reads were used to polish the resulting assemblies using POLCA (from MaSuRCA version 3.4.2) (Zimin et al. 2013). Genome assemblies were visualized with Bandage (Wick et al. 2015). The resulting original genome assemblies were separately annotated with MAKER (version 2.31.10)(Campbell 2014) for both species, using an iterative approach to train AUGUSTUS (version 3.3.3)(Stanke 2006) and SNAP (version 2006.07.28-1)(Korf 2004) gene-model predictors; initial input for all annotation pipelines included the set of transcript and protein sequences from the published *N. thermarum* genome assembly/annotation(Povilus 2020), the set of all protein sequences from Nymphaeaceae available on NCBI, protein sequences from the *N. colorata* genome assembly and annotation (Zhang 2020), all basal Magnoliophyta protein sequences on Uniprot, Amborella protein sequences, and TAIR10 protein sequences. Three rounds of annotation and gene model predictor training were performed for annotation of both species. Repeat identification and masking was performed with RepeatMasker (version 4.0.5)(Chen 2004) using Spermatophyta as the specified query clade and the Embryophyta repeat database.

To create genome assemblies of *N. thermarum* and *N. dimorpha* that shared positional homology, the *N. thermarum* contigs were mapped to *N. dimorpha* contigs using minimap2 (Li 2018) (the *N. dimorpha* assembly was used as the reference as it had the fewest contigs; genome alignment visualized using D-Genies (Cabanettes and Klopp 2018). The resulting reorganized genomes for both species were separately re-annotated as described above, and the final annotations were resolved using MAKER to the reorganized *N. thermarum* genome, with positional homology used to apply the annotation to the *N. dimorpha* genome assembly. For each species, assembled transcripts were then generated using the resolved annotation and genome assembly of each species, resulting in a set of homologous *N. thermarum* transcripts and a set of *N. dimorpha* transcripts.

For *N. thermarum* and *N. dimorpha* transcripts, homology to *Arabidopsis thaliana* was determined by blastx searching the *N. thermarum* transcripts against the TAIR11 protein set (e-value cut-off set at 1e-4). The top *A. thaliana* blastx hit for each *N. thermarum* transcript was selected as the putative homolog. For each *N. dimorpha* transcript, the putative *N. thermarum* homolog was similarly identified with a blastx search (e-value cut-off set at 1e-4), and the corresponding *A. thaliana* homologs was assigned to the *N. dimorpha* transcript.

### DNA methylation-sensitive sequencing and analysis

DNA was extracted from endosperm using the QIAamp micro kit (Qiagen, cat# 56304). For enzymatic methyl conversion sequencing and library preparation, an NEBNext Enzymatic Methyl-seq kit was used; one additional AMPure bead clean-up was performed on libraries to remove primer dimer. Sequencing was performed at the Whitehead Institute Genome Technology Core. Libraries were pooled and sequenced across 2 lanes of a NovaSeq SP flowcell (50 bp, paired-end reads) (endosperm samples) or 2 lanes of a NovaSeq S4 flowcell (150 bp, paired-end) (leaf samples) to give ∼14x genome coverage. Enzymatic-methyl sequencing conversion rate was assessed prior to sequencing (Supplementary Materials and Methods). Conversion rates were calculated using CyMATE (Hetzl et al. 2007). Sample conversion rate averaged 99.85%.

Reads from enzymatic-converted samples were first mapped to a concatenation of the originally produced *N. thermarum* and *N. dimorpha* assemblies and annotations, using Bismark (version 0.22.3) (Krueger and Andrews 2011). 150 bp reads of leaf samples were broken into 40 bp segments and all reads were treated as single-end during mapping to ensure consistency in data processing. The reads that uniquely mapped to either species’ genome were sorted into separate sets of *N. thermarum* or *N. dimorpha* reads and mapped to their respective species’ reorganized genome annotation with Bismark and methylation data was extracted. Analysis of average DNA methylation 5’, 3’ and interior of features was performed using previously developed custom pipelines (Pignatta et al. 2014). Differentially methylated regions (DMRs) between samples were identified in the CG, CHG, and CHH contexts using a previously developed pipeline (Pignatta et al. 2014). DMRs were defined as 300-bp windows for which 3 or more cytosines with a coverage of 5 or more reads had a methylation difference of 35% or greater between samples for CG and CHG contexts and 10% or greater for the CHH context, with a Fisher’s exact test with Benjamini-Hochberg correction p-value cutoff of 0.01 to determine significance. DMRs were called between all combinations of biological replicates. For total number of DMRs between endosperm maternal and paternal alleles, the number of DMRs was averaged across all replicate comparisons. Genes and repeat regions were identified as associated with a DMR if the gene or repeat region had a DMR within the annotated region or +/-2 kb.

GO enrichment analysis was performed using ShinyGO 0.77 (Ge et al. 202). Putative Arabidopsis homologs of all transcripts were used, and the set of putative Arabidopsis homologs of all transcripts expressed during seed development (TPM > 1) (Povilus and Friedman 2022) was used as the background set.

### RNA sequencing and data analysis

For mRNA sequencing, RNA was extracted from frozen endosperm samples via RNAqueous Total RNA Isolation Kit (Invitrogen) according to the kit protocol. Libraries were prepared and sequenced at the MIT BioMicroCenter via NEBNext Ultra II Directional RNA Library Prep Kit for Illumina (polyA-based isolation). Samples were pooled and sequenced on one NovaSeq S4 flowcell (50bp, single end reads).

For full analysis methods, see Supplementary Materials and Methods. Briefly, for initial analysis of gene expression, reads from all <u>hybrid</u> samples were mapped to the concatenated genomes of the originally produced *N. thermarum* and *N. dimorpha* assemblies and annotations<u>; reads from non-hybrid samples were mapped to the reorganized genome of their respective species</u>. For identification of imprinted genes in hybrid samples, the reads that uniquely mapped to either species’ genome were sorted into separate sets of *N. thermarum* or *N. dimorpha* reads and were used for subsequent analysis. *N. thermarum* reads were mapped to the reorganized *N. thermarum* genome annotation, *N. dimorpha* reads were mapped to the reorganized *N. dimorpha* genome annotation. Resulting allele-specific count tables for each transcript were used for calling genetic imprinting. Genetically imprinted genes were called as previously described (Gehring et al. 2011; Pignatta et al. 2014), using a pairwise comparison of all possible combinations of each hybrid cross sample. For each gene we tested whether there was a significant difference (Benjamini-Hochberg adjusted p-value < 0.01) between *p_1_* and *p_2_*, where *p_1_* is the portion of Nt reads for a gene in a Nt x Nd cross and *p_2_* is the portion of Nt reads for the same gene in a Nd x Nt cross. While mapping reads, a slight maternal expression bias was noted for both cross directions (Supplemental Table 1). Therefore, when calling imprinted genes, the expected maternal : paternal expression ratio was adjusted from 1 (the anticipated null ratio for diploid endosperm) to the average maximum observed maternal expression bias of 1.32 (null hypothesis: *p_1_*=1.32, *p_2_*=0.57). To increase stringency, minimum allelic-specific read count was set to 50, a minimum imprinting factor was set to 2, and a maximum cis-effect factor was set to 15. The imprinting factor is a measure of the magnitude of imprinting. For each gene in a sample, a 95% confidence interval was determined around the Nt/Nd read ratio; the imprinting factor is the low value of the high confidence interval divided by the high value of the low confidence interval for the reciprocal cross (Gehring et al. 2011). The *cis*-effect factor is calculated in a similar manner. In addition to these specifications, MEGs were required to have a minimum of 70% maternal allele reads and PEGs were required to a maximum of 30% maternal allele reads in both cross directions. In order for a gene to be considered as consistently imprinted, it had to be called as imprinted in at least 75% (3 of 4) of pairwise comparisons.

For correction of endosperm reads to account for potential maternal tissue contamination, we mapped reads from endosperm samples (this study) and whole-seed samples (Povilus and Friedman 2022) to the reorganized *N. thermarum* genome and proceeded as described in (Tonosaki et al. 2024) (Supplementary Materials and Methods).

Differential gene expression analysis between endosperm of hybrid crosses and non-hybrid endosperm was performed by mapping reads to the concatenated genomes as described above, and then mapping uniquely mapping reads to their respective reorganized genome using kallisto (v 0.46.1) (Bray et al. 2016). Differential gene expression analysis was performed using DEseq2 (Love et al. 2014) using mostly default settings and filtering for loci with adjusted p value less than or equal to 0.01 and mean TPM (of all samples) greater than or equal to 10.

GO enrichment analysis was performed using ShinyGO 0.77 (Ge et al. 2020), using default settings. Putative Arabidopsis homologs of all transcripts in the test set were used, and the set of putative Arabidopsis homologs of all transcripts expressed during seed development (TPM > 1) (Povilus and Friedman 2022) was used as the background set.

### In situ hybridizations and histology

*In situ* hybridizations were performed as previously described (Pignatta et al. 2018) (see Supplementary Materials and Methods for probe information). Preparation of seed samples for histological analysis was performed as previously described for seeds of *A. thaliana* (Pignatta et al. 2018) and stained with toluidine blue, with adaptations of incubation times as necessary. All samples were sectioned on a Leica RM 2065 rotary microtome at a thickness of 8 μm and imaged using a Zeiss Axio Imager M2. Image tiling, color and brightness/contrast adjustments and Smart Sharpen were applied to whole images, with particular attention to having even contrast and white-balance across different images (Adobe Photoshop).

## Acknowledgments

Funding was provided by the National Science Foundation grants MCB 2101337 to M. G., MCB 1453459 to M. G., IOS 1812116 to R.A.P, and a NSF Graduate Research Fellowship to C.A.M. Sequencing was performed at the MIT BioMicro Center and the Whitehead Institute Genome Technology Core.

## Notes

https://www.ncbi.nlm.nih.gov/bioproject/PRJNA1085992

https://www.ncbi.nlm.nih.gov/bioproject/PRJNA1085993

https://www.ncbi.nlm.nih.gov/bioproject/PRJNA1086863

https://www.ncbi.nlm.nih.gov/bioproject/PRJNA1086866

https://www.ncbi.nlm.nih.gov/bioproject/PRJNA1087317

### Competing Interest Statement

The authors have declared no competing interest.

### Summary of Updates

additional dataset and analysis with associated results and discussion, as well as additional details of materials and methods and updated references cited.

