## supplemental information for "Imprinting and DNA methylation in water lily endosperm: implications for seed evolution"

For:

### PDF Contents:

Supplementary Figures, 1-5

Supplementary Table Titles, 1-5

Supplementary Materials and Methods

### Supplementary Figures

**Supplementary Figure 1: Image of individuals of *N. thermarm*, *N. dimorpha*, and an F1 hybrid.**
Top left = *N. thermarum*, top right = *N. dimorpha*, bottom = F1 hybrid (demonstrating viability of hybrid crosses). Identity of individuals was confirmed by genotyping (EM seq samples s104, 105, s106, see Supplementary Table 1).

Supplementary Figure 1

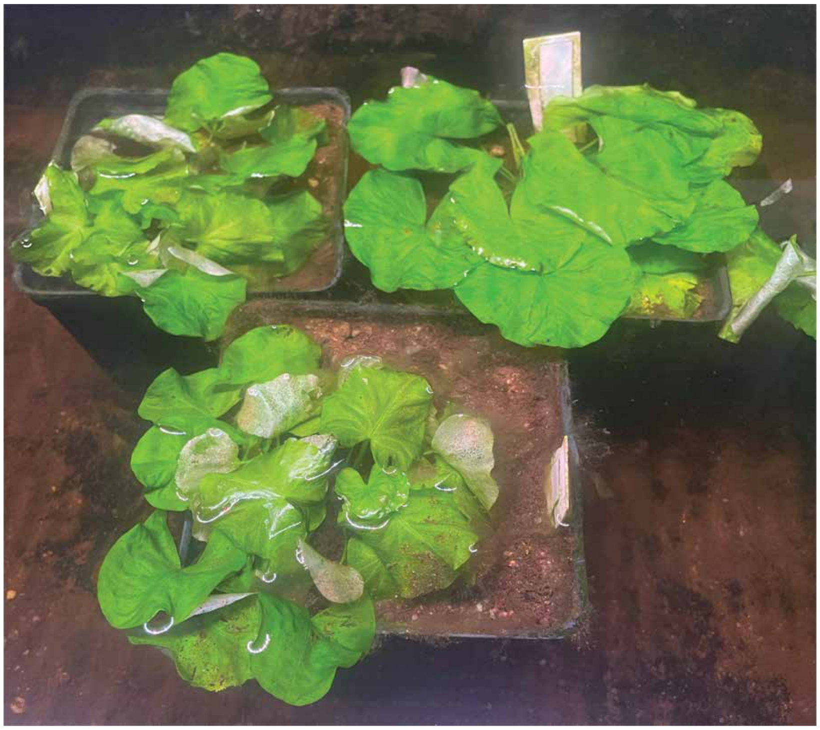

**Supplementary Figure 2: Genome assemblies for *N. thermarum* and *N. dimorpha*.** A) Bandage diagrams for the updated *N. thermarum* genome assembly and the novel *N. dimorpha* genome assembly. B) Alignment of the *N. thermarum* genome mapped against the *N. dimorpha* genome.

Supplementary Figure 2

A

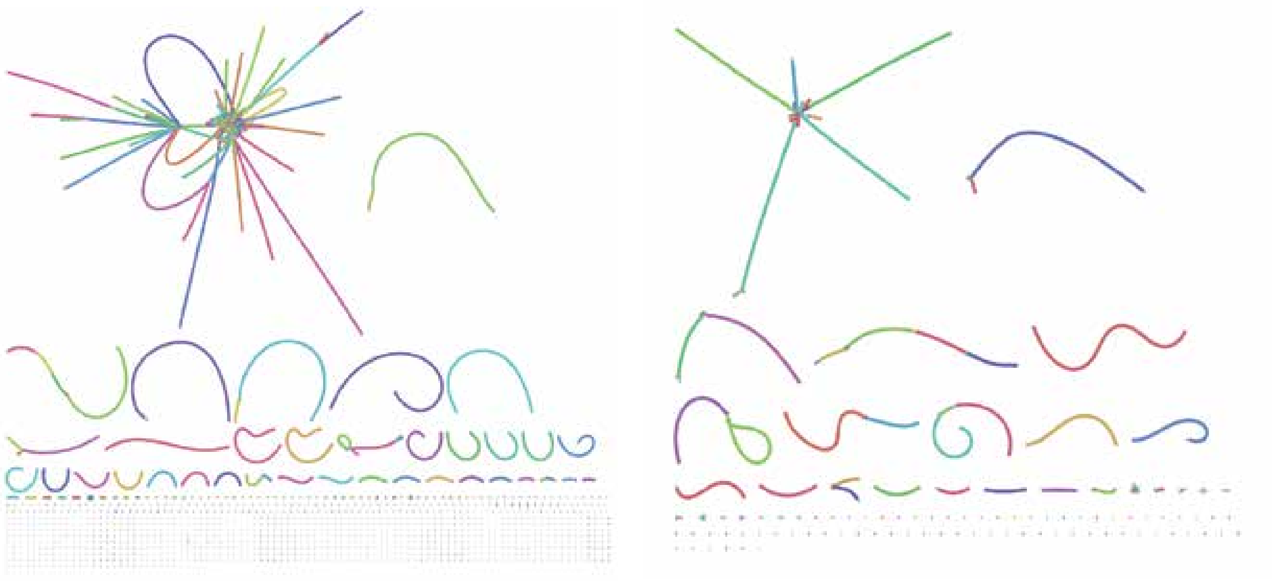

*Nymphaea thermarum*
1,553 contigs

*Nymphaea dimorpha*
248 contigs

B

*Nymphaea dimorpha* genome
("nymmin" and "nm" refer to *N. dimorpha*. *N. dimorpha* replaced the taxon synonym *N. minuta*)

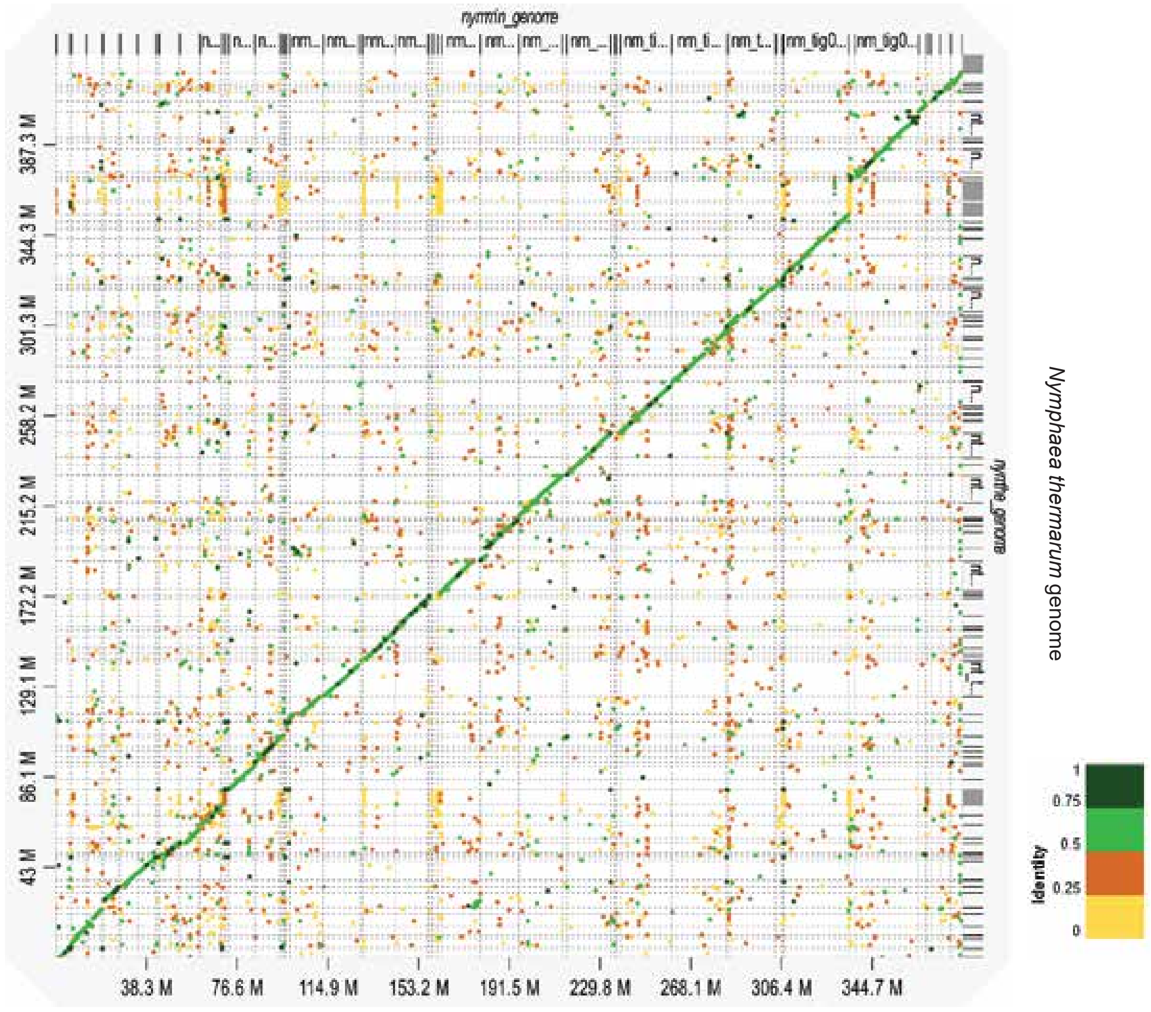

**Supplementary Figure 3: Differential gene expression analysis between hybrid and parental endosperm with MEGs highlighted.** A) Comparison of fold change and averaged expression (TPM) for *N. dimorpha* self-fertilized endosperm vs. *N. thermarm* x *N. dimorpha* hybrid endosperm. B) Comparison of fold change and averaged expression (TPM) for *N. dimorpha* self-fertilized endosperm vs. *N. dimorpha* x *N. thermarum* hybrid endosperm. C) Comparison of fold change and averaged expression (TPM) for *N. thermarum* self-fertilized endosperm vs. *N. thermarm* x *N dimorpha* hybrid endosperm. D) Comparison of fold change and averaged expression (TPM) for *N. thermarum* self-fertilized endosperm vs. *N. dimorpha* x *N. thermarum* hybrid endosperm. E) Upset graph showing overlap of MEGs that are significantly DE for the the four different comparison types; the putative *Arabidopsis thaliana* homologs of the 10 consistently DE MEGs is listed.

Supplementary Figure 3

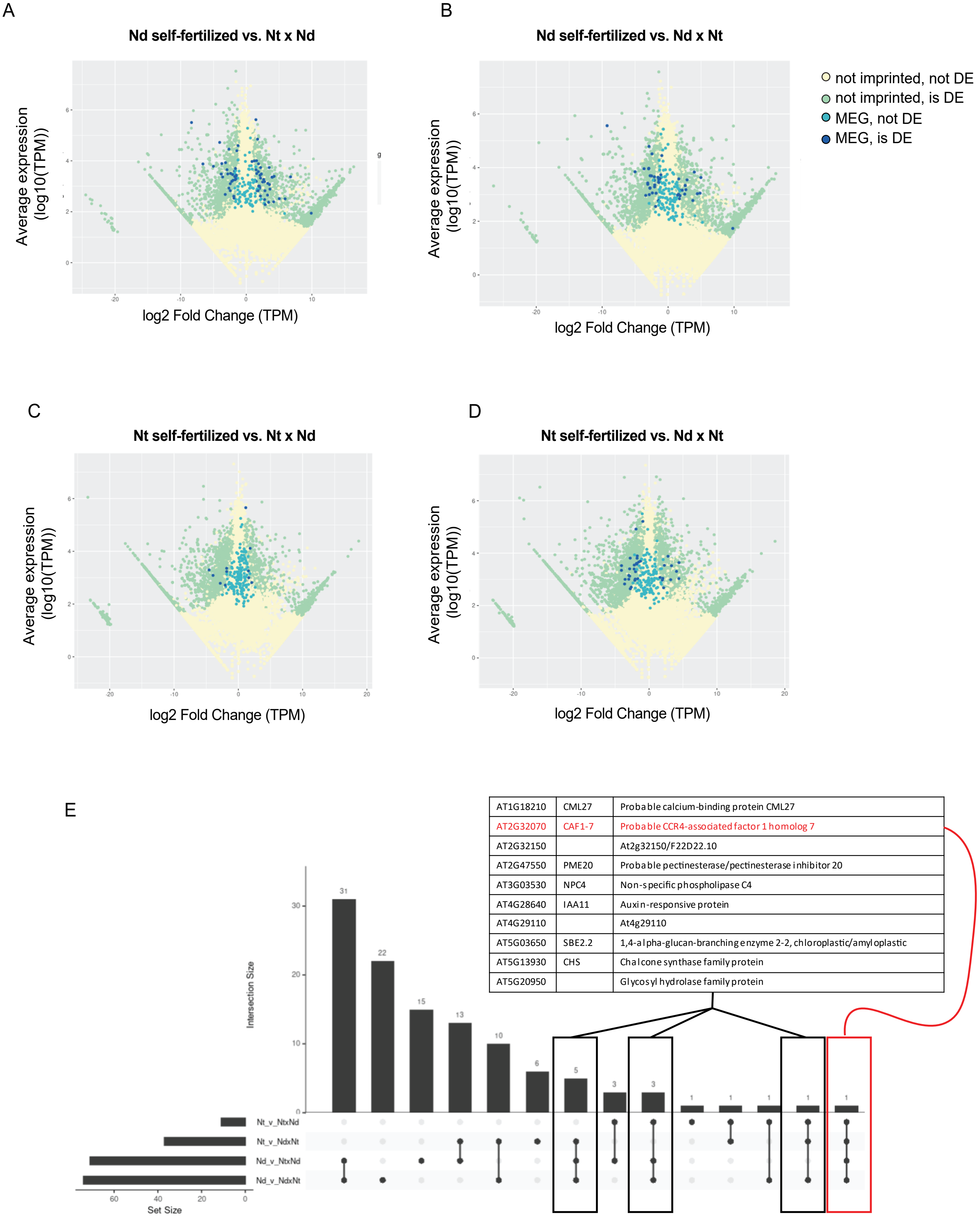

**Supplementary Figure 4: In situ hybridization positive control for perisperm, and SUS3 expression in seeds of different crosses and self-fertilizations.** A) RNA in situ hybridization of putative homolog of a terpene synthase sub-family in *N. thermarum* seeds, showing that detection of gene expression in the perisperm is possible. Results of experiments performed with antisense and sense probes are shown. Scalebars = 200 µm. B) In situ hybridization of putative homolog of *SUS3*, performed in seeds of reciprocal crosses of *N. thermarum* and *N. dimorpha* and in seeds from *N. dimorpha* self-fertilizations. Black arrowheads indicate detection of signal, white arrowheads indicate absence of signal. Scalebars = 200 µm. Results of experiments performed with antisense and sense probes are shown.

Supplemental Figure 4

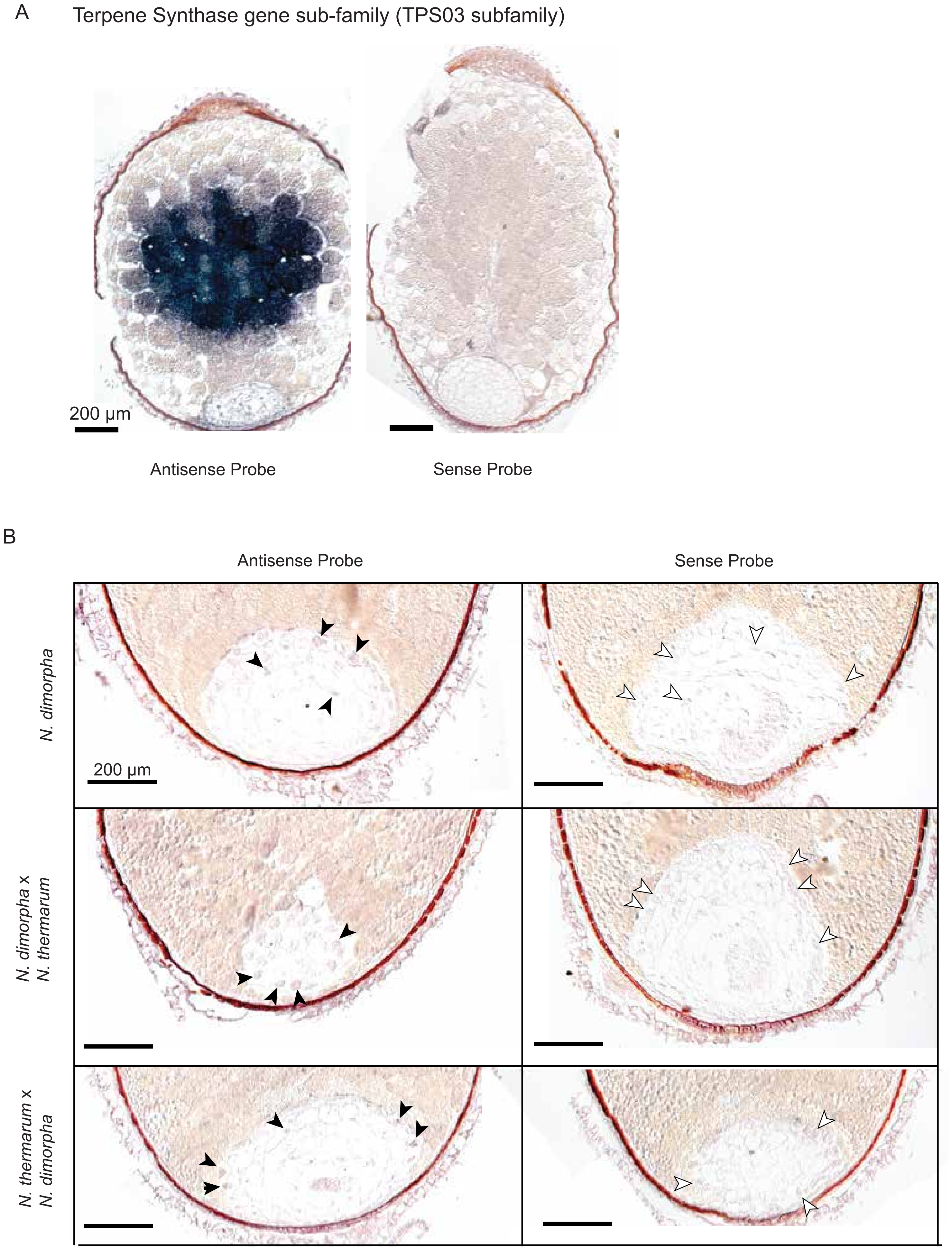

**Supplementary Figure 5: DNA methylation, including leaf tissues from non-hybrid *N, thermarum* and *N. dimorpha* plants**

A) Number of DMRs that are hyper- or hypomethylated in the *N. dimorpha* compared to *N. thermarum* genome, that overlap a gene and/or a repeat region, for each species, in leaf tissue. DMRs were called by comparing the genomes of each species within the same sample (hybrid tissue, one sample) or between samples of leaves of each parental species (one sample of each species). B) Genome browser snapshots of DNA methylation for homologs of *GA2OX2* and *TMO6*, showing examples of methylation patterning (blue = CG, red = CHG, and green = CHH) on the genome for each species , in endosperm and leaf tissue, including leaves of non-hybrid *N, thermarum* and *N. dimorpha* plants. Black notches indicate cytosines for which there was sufficient data to include, but were unmethylated.

Supplementary Figure 5

A

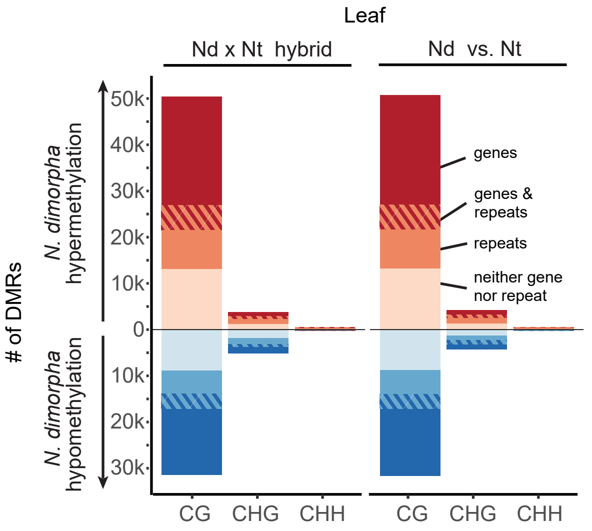


B

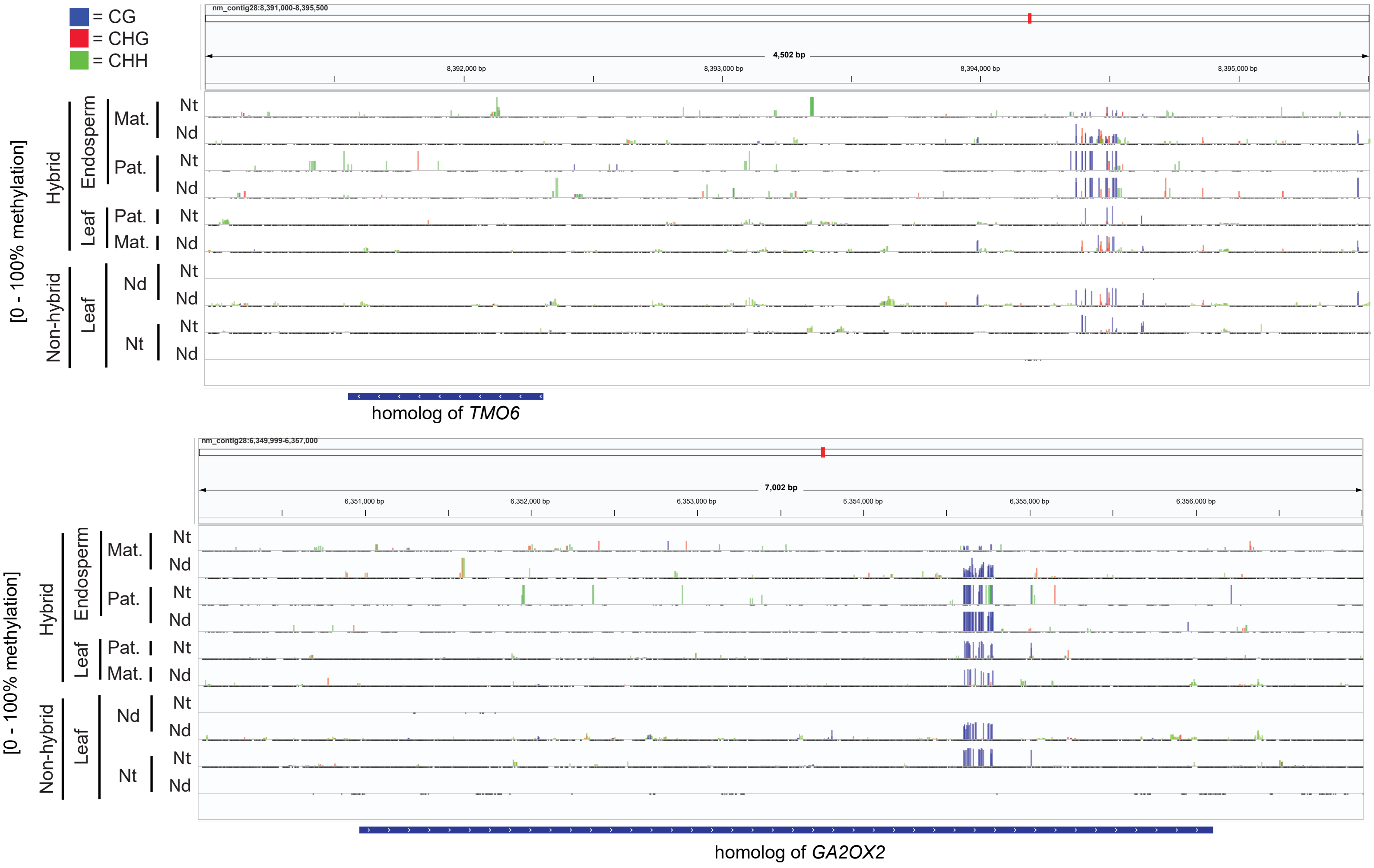

### Supplementary Tables, Titles

**Supplementary Table 1:** Summaries for sequencing, mapping, and genome assemblies and annotations.

**Supplementary Table 2:** Results and summaries of imprinting calling tests, report of imprinted loci that overlap DMRs, and summary of imprinting status of *Nymphaea* homologs of conserved imprinted genes.

**Supplementary Table 3:** Results and summaries of differential gene expression analysis between intraspecies and interspecies crosses

**Supplementary Table 4:** Results and summaries of imprinting calling tests, when data is adjusted for potential maternal tissue contamination of endosperm samples (assuming 50% and 25% contamination).

**Supplementary Table 5**: Report of DMR overlap with genomic features, as well as report and summary of genes with DMR overlaps

### Supplementary Materials and Methods

*(additional Materials and Methods details)*

### Plant growth and sample collection

Seeds of *N. thermarum* and *N. dimorpha* were sourced from the Arnold Arboretum of Harvard University (Boston, MA, USA) and grown at the Whitehead Institute for Biomedical Research (Cambridge, MA, USA). Plants were grown in individual pots, potted in top soil, and submerged in pH-balanced tap water (pH roughly between 6.5 and 7). *N. thermarum* plants were submerged so that the soil level was 2-5 cm below the water level, while *N. dimorpha* plants were submerged so that the soil level was 15-30 cm below the water level. Plants were grown under a mix of natural and artificial lighting (full spectrum LED), with seasonal lighting supplemented to provide 12 h of light a day. To encourage prolific flowering and fruit set, every 2-4 weeks approximately one half of a CrystalClear Thrive 10-14-8 Aquatic Plant Fertilizer tablet was added to each pot.

Controlled pollinations and self-fertilizations were performed as previously described [Povilus 2015 ][Povilus 2018]. For collection of seeds from crosses and self-fertilizations, first day of anthesis (time of female receptivity and fertilization [Povilus 2015]) was defined by presence of stigmatic fluid. Fruits were collected at 9-10 days after anthesis (DAA) and seeds were immediately removed and dissected with fine forceps in dissection buffer (0.3M sorbitol, 0.3mM MES) to separate the embryo and endosperm; endosperms were placed in droplets of fresh dissection buffer. For each sample, tissue from 10-20 seeds from a single fruit were pooled. Endosperms were washed by moving to a fresh droplet of dissection buffer via forceps or micropipette; samples were then moved to a third and final droplet of fresh dissection buffer before being moved to a microcentrifuge tube. Excess dissection buffer was drawn off and samples were frozen in liquid nitrogen.

**Whole genome sequencing, assembly, and annotation**

For long-read and short-read DNA sequencing, high molecular weight DNA was extracted from > 1 g young leaf samples from a single individual plant using a modified CTAB-based protocolBriefly: frozen leaf tissue was ground in a mortar and pestle and added to a CTAB buffer with 0.25 mM PVP40 (Polyvinylpyrrolidone), 0.005% (v/v) 2-Mercaptoethanol, and 1.3% (v/v) sodium dodecyl sulfate. Samples were incubated for 30 minutes at 65 °C, centrifuged to precipitate debris, and supernatant was removed. Samples were precipitated with addition of potassium acetate (to 0.9M), incubated on ice for 30 minutes and then centrifuged at 4 °C for 20 min. Supernatant was transferred to a new tube and an equal volume of isopropanol was added; samples were incubated on ice for 5 minutes, centrifuged at 4 °C for 20 minutes, and supernatant was removed and discarded. The air-dried DNA pellet was re-suspended in Tris-EDTA buffer; RNAaseA was added and samples incubated for 20 minutes at 37 °C. Samples were transferred to PhaseLock Heavy Gel tubes, and phase-extracted 3 times with 25:24:1, phenol : chloroform : isoamyl alcohol and then 3 times with chloroform (the aqueous phase was transferred to a new PhaseLock gel tube for each step). Samples were then precipitated with sodium acetate (to 0.3 M) and an equal volume of isopropanol; samples were chilled on ice for 5 min and centrifuged at 4 °C for 1 minute. DNA pellets were washed three times with 70% ethanol, air dried and re-suspended in 0.1M Tris-EDTA buffer.

Genome assembly for *N. thermarum* and *N. dimorpha* was performed separately using long-reads as input for Canu (version 2.1.1) [Koren 2017]; short-reads were used to polish the resulting assemblies using POLCA (from MaSuRCA version 3.4.2) [Zimin 2013]. Genome assemblies were visualized with Bandage[Wick 2015]. The resulting original genome assemblies were separately annotated with MAKER (version 2.31.10) [Campbell 2014] for both species, using an iterative approach to train AUGUSTUS (version 3.3.3) [Stanke 2006] and SNAP (version 2006.07.28-1) [Korf 2004] gene-model predictors; initial input for all annotation pipelines included the set of transcript and protein sequences from the published *N. thermarum* genome assembly/annotation [Povilus 2020], the set of all protein sequences from Nymphaeaceae available on NCBI, protein sequences from the *N. colorata* genome assembly and annotation [Zhang 2020], all basal Magnoliophyta protein sequences on Uniprot, Amborella protein sequences, and TAIR10 protein sequences. Three rounds of annotation and gene model predictor training were performed for annotation of both species. Repeat identification and masking was performed with RepeatMasker (version 4.0.5) [Chen 2004] using Spermatophyta as the specified query clade and the Embryophyta repeat database.

**DNA methylation-sensitive sequencing and analysis**

Reads from enzymatic-converted samples were first mapped to a concatenation of the originally produced *N. thermarum* and *N. dimorpha* assemblies and annotations, using Bismark (version 0.22.3)[Keuger 2011]. The reads that uniquely mapped to either species' genome were sorted into separate sets of *N. thermarum* or *N. dimorpha* reads. *N. thermarum* reads were mapped to the reorganized *N. thermarum* genome annotation and methylation data was extracted with Bismark. *N. dimorpha* reads were mapped to the reorganized *N. dimorpha* genome annotation and methylation data was extracted with Bismark. Analysis of average DNA methylation 5', 3' and interior of features was performed using previously developed custom pipelines [Pignatta 2014]. Differentially methylated regions (DMRs) between samples were identified in the CG, CHG, and CHH contexts using a previously developed pipeline [Pignatta 2014]. Briefly, DMRs were defined as 300-bp windows for which 3 or more cytosines with a coverage of 5 or more reads had a methylation difference of 35% or greater between samples for CG and CHG contexts and 10% or greater for the CHH context, with a Fisher's exact test with Benjamini-Hochberg correction p-value cutoff of 0.01 to determine significance. DMRs were called between all combinations of biological replicates. For total number of DMRs between endosperm maternal and paternal alleles, the number of DMRs was averaged across all replicate comparisons. Genes and repeat regions were identified as associated with a DMR if the gene or repeat region had a DMR within the annotated region or +/-2 kb.

**RNA sequencing and data analysis**

For initial analysis of gene expression, reads from all hybrid samples were mapped to the concatenated genomes of the originally produced *N. thermarum* and *N. dimorpha* assemblies and annotations, using STAR (version 2.4.1) using default settings, after using TrimGalore (version 0.6.7) to trim reads of adaptors and low-quality regions. Reads from non-hybrid samples were trimmed using TrimGalore and mapped to the reorganized genome of their respective species via Kallisto (version 0.46.1). For identification of imprinted genes in hybrid samples, the reads that uniquely mapped to either species' genome were sorted into separate sets of *N. thermarum* or *N. dimorpha* reads, and were used for subsequent analysis. *N. thermarum* reads were mapped to the reorganized *N. thermarum* genome annotation via Kallisto (version 0.46.1), using the set of *N. thermarum* transcripts generated from the reorganized genome assembly and annotation. *N. dimorpha* reads were mapped to the reorganized *N. dimorpha* genome annotation via Kallisto, using the set of *N. dimorpha* transcripts generated from the reorganized genome assembly and annotation. Resulting allele-specific count tables for each transcript were used for calling genetic imprinting. Genetically imprinted genes were called as previously described [Pignatta 2014], using a pairwise comparison of all possible combinations of each hybrid cross sample (*N. thermarum* x *N. dimorpha* vs. *N. dimorpha* x *N. thermarum*). While mapping reads, a slight maternal expression bias was noted for both cross directions [Supplemental Table 1]. Therefore, when calling imprinted genes, the expected maternal : paternal expression ratio was adjusted from 1 (the anticipated null ratio for diploid endosperm) to the average maximum observed maternal expression bias of 1.32; to increase stringency minimum allelic-specific read count was set to 50, a minimum imprinting factor was set to 2, and cis-effect factor was set to 15. MEGs were required to have a minimum of 70% maternal allele reads and PEGs were required to a maximum of 30% maternal allele reads in both cross directions. In order for a gene to be considered as consistently imprinted, it had to be called as imprinted in at least 75% (3 of 4) of pairwise comparisons. For total read counts from hybrid samples, reads that mapped twice to the concatenated genomes (mapq score =3) were filtered and mapped to one reorganized genome (*N. dimorpha*) via kallisto. The resulting read counts per locus were added to the totals of the read counts that mapped uniquely to the maternal or paternal genomes. Principal components analysis of read counts was performed in R using the DESeq2 package [Love 2014], including using the varianceStabilizingTransformation() function.

For correction of endosperm reads to account for potential maternal tissue contamination, we mapped reads from endosperm samples (this study) and whole-seed samples [Povilus 2022a] to the reorganized *N. thermarum* genome and proceeded as described in [Tonosaki 2023]. For each expressed transcript we used the upper bound on the 95% binomial confidence intervals of the ratio of averaged total endosperm read counts (in TPM, reads mapping to both the maternal and paternal genomes) to averaged whole-seed transcript counts (in TPM), assumed that either 50% or 25% of the endosperm transcript pool consisted of transcripts from the whole-seed, and calculated the resulting "real endosperm" read counts mapping to the maternal genome before calling imprinted genes (parental genome specific mapping and imprinting calling as described above). This correction was only done for maternal read counts.

**In situ hybridizations and histology**

In situ hybridizations were performed as previously described[Pignatta 2018] [Supplementary Materials and Methods]. Probe primer sequences were as follows: GA2OX2, forward = 5' ATCTAACCCAGCACTCGACC 3' ; GA2OX2, reverse = 5 'CTGTGGTGAAGGTGTCAAGC 3'; SUS3, forward = 5' GCTACTTTATTGGCACATAAGCTG 3'; SUS3, reverse = 5' GAAATCGCATCCCAGTGTTG 3'; Terpene synthase gene subfamily, forward = 5' CTTGTGAGATGTGAGCCTAGG 3'; Terpene synthase gene subfamily, reverse = 5' TGCCACCTTGCCTCAAC 3'. Anticipated probe lengths were as follows: GA2OX2 = ~900 bp; SUS3 = ~1000 bp; terpene synthase gene subfamily = ~1000 bp.

**References, Supplementary Materials and Methods**

[Campbell 2014] M. S. Campbell, C. Holt, B. Moore, M. Yandell, Genome Annotation and Curation Using MAKER and MAKER-P. *Curr. Protoc. Bioinformatics* 48, 4.11.1–4.11.39 (2014). 10.1002/0471250953.bi0411s48.

[Chen 2004] N. Chen, Using RepeatMasker to identify repetitive elements in genomic sequences. *Curr. Protoc. Bioinformatics*, Chapter 4 (2004). 1002/0471250953.bi0410s05.

[Koren 2017] S. Koren, B. P. Walenz, K. Berlin, J. R. Miller, A. M. Phillippy, Canu: scalable and accurate long-read assembly via adaptive k-mer weighting and repeat separation. *Genome Res.* 27, 722-736 (2017). 10.1101/gr.215087.116

[Korf 2004] I. Korf, Gene finding in novel genomes. *BMC Bioinform.* 5, 59 (2004). 10.1186/1471-2105-5-59.

[Love 2014] M. I. Love, W. Huber, S. Anders, Moderated estimation of fold change and dispersion for RNA-seq data with DESeq2. *Genom. Biol.* 15, 550 (2014). doi:10.1186/s13059-014-0550-8.

[Pigantta 2014] D. Pignatta, R. M. Erdmann, E. Scheer, C. L. Picard, G. W. Bell, M. Gehring, Natural epigenetic polymorphisms lead to intraspecific variation in Arabidopsis gene imprinting. *Elife*, 3:3:e03198 (2014). 10.7554/eLife.03198.

[Povilus 2015] R. A. Povilus, J. M. Losada,W. E. Friedman, Floral biology and ovule and seed ontogeny of Nymphaea thermarum, a water lily at the brink of extinction with potential as a model system for basal angiosperms. *Ann. Bot.* 115, 211–226 (2015). 10.1093/aob/mcu235.

[Povilus 2018] R. A. Povilus, P. K. Diggle W. E. Friedman, Evidence for parent-of-origin effects and interparental conflict in seeds of an ancient flowering plant lineage. *Proc. Royal. Soc. B* 285(1872): 20172491 (2018). 10.1098/rspb.2017.2491

[Povilus 2020] R. A. Povilus, J. M. DaCosta, C. Grassa, P. R. V. Satyaki, M. Moeglein, J. Jaenisch, Z. Xi, S. Mathews, M. Gehring, C. C. Davis, W. E. Friedman, Water lily (Nymphaea thermarum) genome reveals viaraible genomic signatures of ancient vascular cambium losses. *Proc. Natl. Acad. Sci. U S A* 117(15), 8649-8656 (2020). doi.org/10.1073/pnas.1922873117

[Povilus 2022a] R. A. Povilus, W. E. Friedman, Transcriptomes across fertilization and seed development in the water lily Nymphaea thermarum: evidence for epigenetic patterning during reproduction. *Plant Reprod.* 35(5), 161-178 (2020). 10.1007/s00497-022-00438-3.

[Stanke 2006] M. Stanke, O. Keller, I. Gunduz, A. Hayes, S. Waack, B. Morgenstern, AUGUSTUS: ab initio prediction of alternative transcripts. *Nucleic Acids Res.* 34, W435–W439 (2006). 10.1093/nar/gkl200.

[Tonosaki 2023] K. Tonosaki, A. Ono, H. Nagata, H. Furuumi, K. Nonomura, Y. Sato, L. Comai, K. Hatakeyama, T. Kamakatsu, T. Kinoshita, Multi-layered epigenetic control of persistenet and stage-specific imprinted genes in rice endosperm. *Nat. Plants* 10, 1231–1245 (2024), 10.1038/s41477-024-01754-4.

[Zhang 2020] L. Zhang, F. Chen, X. Zhang, Z. Li, Y. Zhao, R. Lohaus, X. Chang, W. Dong, S. Y. W. Ho, X. Liu, A. Song, J. Chen, W. Guo, Z. Wang, Y. Zhuang, H. Wang, X. Chen, J. Hu, Y. Liu,

Y. Qin, K. Wang, S. Dong, Y. Liu, S. Zhang, X. Yu, Q. Wu, L. Wang, X. Yan, Y. Jiao, H. Kong, X. Zhou, C. Yu, Y. Chen, F. Li, J. Wang, W. Chen, X. Chen, Q. Jia, C. Zhang, H. Ma, Y. Van de Peer, H.Tang, The water lily genome and the early evolution of flowering plants. *Nature* 57(7788), 79-84 (2020). 10.1038/s41586-019-1852-5.

[Zimin 2013] A. V. Zimin, G. Marçais, D. Puiu, M. Roberts, S. L. Salzberg, J. A. Yorke, The MaSuRCA genome assembler. *Bioinformatics* 29(21), 2669-77 (2013). 10.1093/bioinformatics/btt476.
